# Cell autonomous regulation of the activation of AgRP neurons by the melanocortin-3 receptor

**DOI:** 10.1101/2023.06.28.546874

**Authors:** Yijun Gui, Naima S. Dahir, Griffin Downing, Patrick Sweeney, Roger D. Cone

**Affiliations:** Life Sciences Institute, University of Michigan, Ann Arbor, MI; Department of Molecular, Cellular, and Developmental Biology, University of Michigan, Ann Arbor, MI; Department of Molecular and Integrative Physiology, University of Michigan, Ann Arbor, MI; Department of Molecular and Integrative Physiology, University of Illinois, Urbana-Champaign, IL

## Abstract

The melanocortin-3 receptor (MC3R) is a negative regulator of the central melanocortin circuitry via presynaptic expression on AgRP nerve terminals, from where it regulates GABA release onto secondary MC4R-expressing neurons. Hence, animals lacking MC3R (MC3R KO) exhibit hypersensitivity to MC4R agonists. However, MC3R KO mice also exhibit defective behavioral and neuroendocrine responses to fasting. Here, we demonstrate that MC3R KO mice exhibit defective activation of AgRP neurons in response to fasting and cold exposure, while exhibiting normal inhibition of AgRP neurons by sensory detection of food. Further, using an AgRP-specific MC3R knockout model, we show that the control of AgRP neuron activation by MC3R is cell-autonomous. One mechanism underlying this involves the response to ghrelin, which is also blunted in mice with AgRP-specific deletion of the MC3R. Thus, MC3R is a crucial player in the control of energy homeostasis by the central melanocortin system, not only acting presynaptically on AgRP neurons, but via AgRP cell-autonomous regulation of fasting- and cold-induced neuronal activation as well.

## INTRODUCTION

The central melanocortin system, consisting of the primary melanocortinergic AgRP and POMC neurons in the arcuate nucleus and the downstream target sites expressing the melanocortin-3 or -4 receptors (MC3R/MC4R), is critical for the maintenance of energy homeostasis.^1^ The mechanistic link between MC4R deficiency and obesity is well-established,^2-4^ with the MC4R being critical for the establishment of adipostasis, in part via its role in communicating the leptin signal to the CNS.^5^ In contrast, MC3R deficiency in the mouse causes no hyperphagia and a mild, late onset obesity.^6, 7^ One major mechanism functionally distinguishing MC3R from MC4R lies in their distinct expression patterns in the brain,^8, 9^ with *Mc3r* but not *Mc4r*, exhibiting dense mRNA expression in the AgRP and POMC neurons.^10^ The MC3R was discovered to be expressed presynaptically on AgRP neurons, where it regulates GABA release from these terminals onto downstream sites expressing MC3 and/or MC4 receptors.^11^ Chemogenetic manipulation of the ARC MC3R neurons or the pharmacological manipulation of the central MC3R circuitry bidirectionally regulates food intake in a manner phenocopying the manipulation of AgRP neurons.^12^ Further, it has been demonstrated experimentally that the MC3R regulates GABA release onto both POMC neurons^13^ and PVN-MC4R neurons.^11^ Collectively, these data support the model that the MC3R in AgRP terminals may have a global role providing negative regulatory feedback to the entire melanocortin system. Indeed, MC3R KO mice are hypersensitive to both MC4R agonists^11^ and anorexigenic behavioral stimuli^12^, yet also exhibit increased obesity when fed a high fat diet^11^, and this has led to the concept that the MC3R is required for the regulation of the upper and lower boundaries of energy homeostasis, rather than set point itself. However, the MC3R is also required for normal neuroendocrine and behavioral response to fasting,^14, 15^ and the mechanism(s) underlying these observations have not been explained. While MC3R KO mice exhibit normal amounts of ad libitum feeding, a deficit in fast-induced refeeding is readily visible, and persists for many days after a single overnight fast.^11^ MC3R KO mice also exhibit a defect in fasting-induced activation of the hypothalamic-pituitary-adrenal axis,^14^ fasting-induced suppression of the thyroid axis,^12^ and fasting-induced suppression of the hypothalamic-pituitary-gonadal axis.^16^ Absence of the MC3R has also been demonstrated to suppress linear growth and pubertal maturation in both mouse and human.^6, 7, 14, 16^ Here, we identify the cellular basis for the regulation of the fasting response by the MC3R.

## RESULTS

### MC3R is required for the activation of AgRP neurons by fasting

Both male and female MC3R KO mice exhibited defective refeeding in response to a 24-hour fast compared to their wildtype (WT) littermates (Figures 1A-1D), as shown previously.^11, 14^ We next examined hypothalamic mRNA expression by qPCR over a 48-hour fasting time course in both sexes. We found that compared to WT controls, both male and female MC3R KO mice demonstrated significantly impaired fasting-induced upregulation of AgRP and NPY mRNA levels over the entire course of 48h fasting, while the POMC mRNA levels between the two groups were comparable (Figures 1E-1J). Despite relatively reduced expression, there was still a marked upward trend of AgRP and NPY mRNA levels in MC3R KO mice upon fasting, suggesting that the activation of AgRP neurons in MC3R KO mice was suppressed, but not fully abolished.^15^

**Figure 1.**
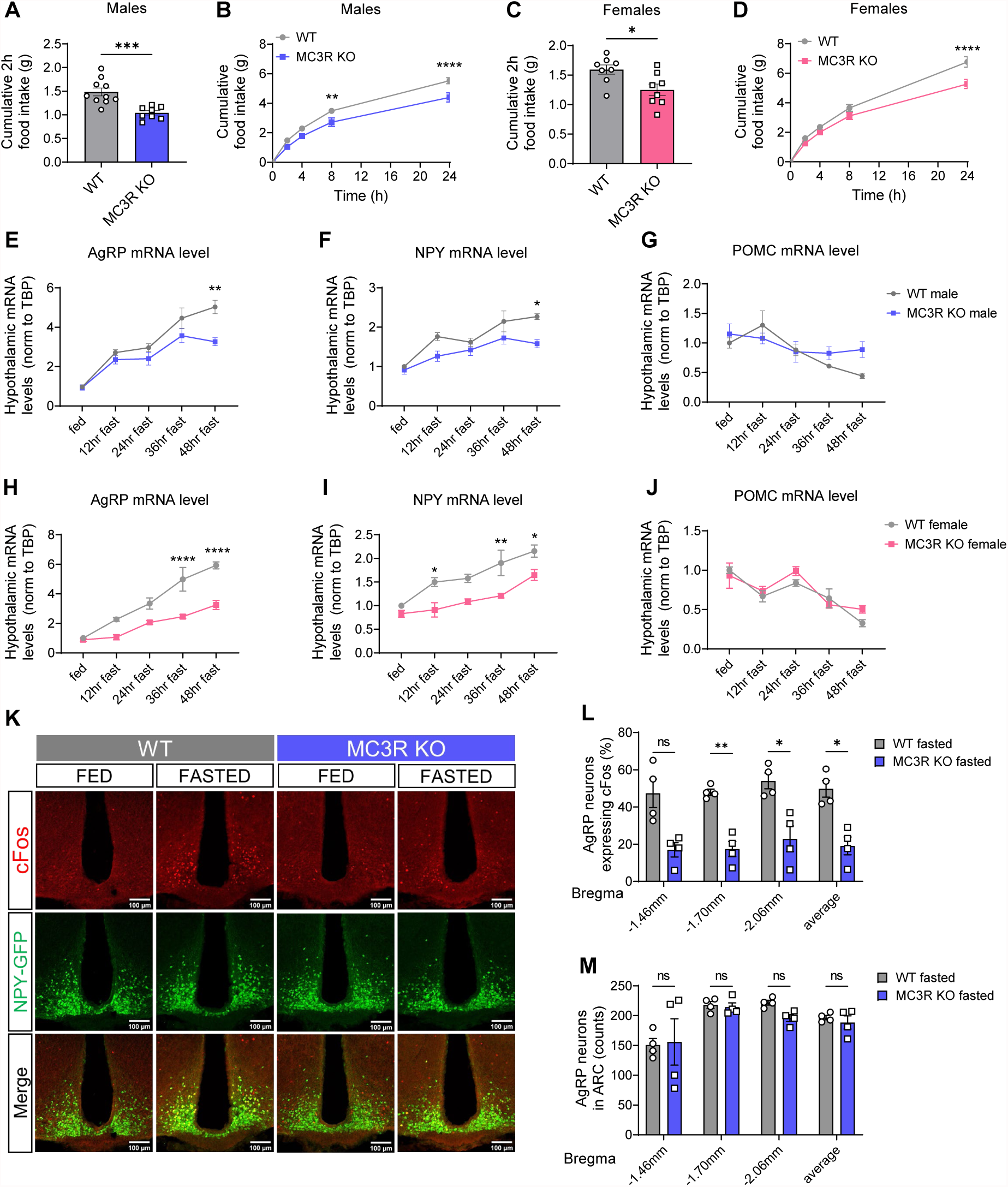
MC3R is required for the activation of AgRP neurons in response to fasting. (A-D) Two-hour (A, C) and 24-hour (B, D) food intake of WT and MC3R KO male and female mice following a 24h fast (n = 8∼10 mice for all groups). (E-G, H-J) qPCR analysis of hypothalamic AGRP, NPY and POMC mRNA levels in WT and MC3R KO male and female mice over a 48h fasting time course (n = 3∼9 mice for all groups). (K-M) Representative images of GFP and cFos immunostaining, and quantifications of the percentage of cFos-positive and the number of GFP cells in the ARC of 24h-fasted WT and MC3R KO NPY-GFP male mice. Scale bar, 100μm (n = 4 mice for all groups). Data are plotted as mean and all error bars represent the SEM. ns, non-significant; *p < 0.05; **p < 0.01; ***p < 0.001, ****p < 0.0001 in unpaired Student’s t-test and two-way ANOVA with Sidak’s posthoc test.

To further investigate the impaired activation of AgRP neurons in MC3R KO mice, we next performed cFos immunostaining in the arcuate nucleus (ARC) of NPY-GFP mice. Compared to the WT group, MC3R KO male and female mice exhibited significantly less cFos expression in response to a 24h fast but not under *ad lib* fed conditions (Figures 1K, 1L, S1A and S1D-E). MC3R KO and WT mice had comparable number of total AgRP neurons regardless of the feeding state (Figures 1M and S1B). Previous work has shown that both AgRP and non-AgRP GABAergic neurons in the ARC can be activated by fasting.^17^ Given the expression of MC3R mRNA detected in non-AgRP cells in the ARC,^12, 18^ we next examined whether fasting activated MC3R-positive non-AgRP neurons in the ARH. While a significant increase in cFos immunoreactivity was seen in non-AgRP MC3R positive ARC cells following fasting, this increase in cFos immunoreactivity was also diminished in the absence of the MC3R (Figure S1C).

### MC3R is required for the activation of AgRP neurons in response to cold

Cold exposure is another example of energy need that evokes adaptive hyperphagia as a compensation for the increased heat production.^19, 20^ While much is known regarding the neurocircuitry underlying cold-induced thermogenesis,^21, 22^ the involvement of AgRP neurons in the regulation of cold-induced hyperphagia had not been discovered until recently.^23, 24^ Given the role of MC3R in regulating AgRP neuron activation by fasting, we tested if MC3R is also required for the cold-induced activation of AgRP neurons. Similarly, we found that both male and female MC3R KO mice exhibited defective hyperphagia compared with WT mice following 4hr cold exposure (4°C versus 22°C) (Figures 2A and 2D). Cold-induced upregulation of AgRP and NPY mRNA levels was largely blunted in MC3R KO mice in both sexes (Figures 2B and 2E). In WT mice, a significant drop in body temperature was detected in response to cold challenge (Figures 2C and 2F) while MC3R KO males but not females showed significantly higher body temperature following cold exposure, indicating increased thermogenesis despite reduced energy intake in the absence of MC3R.

**Figure 2.**
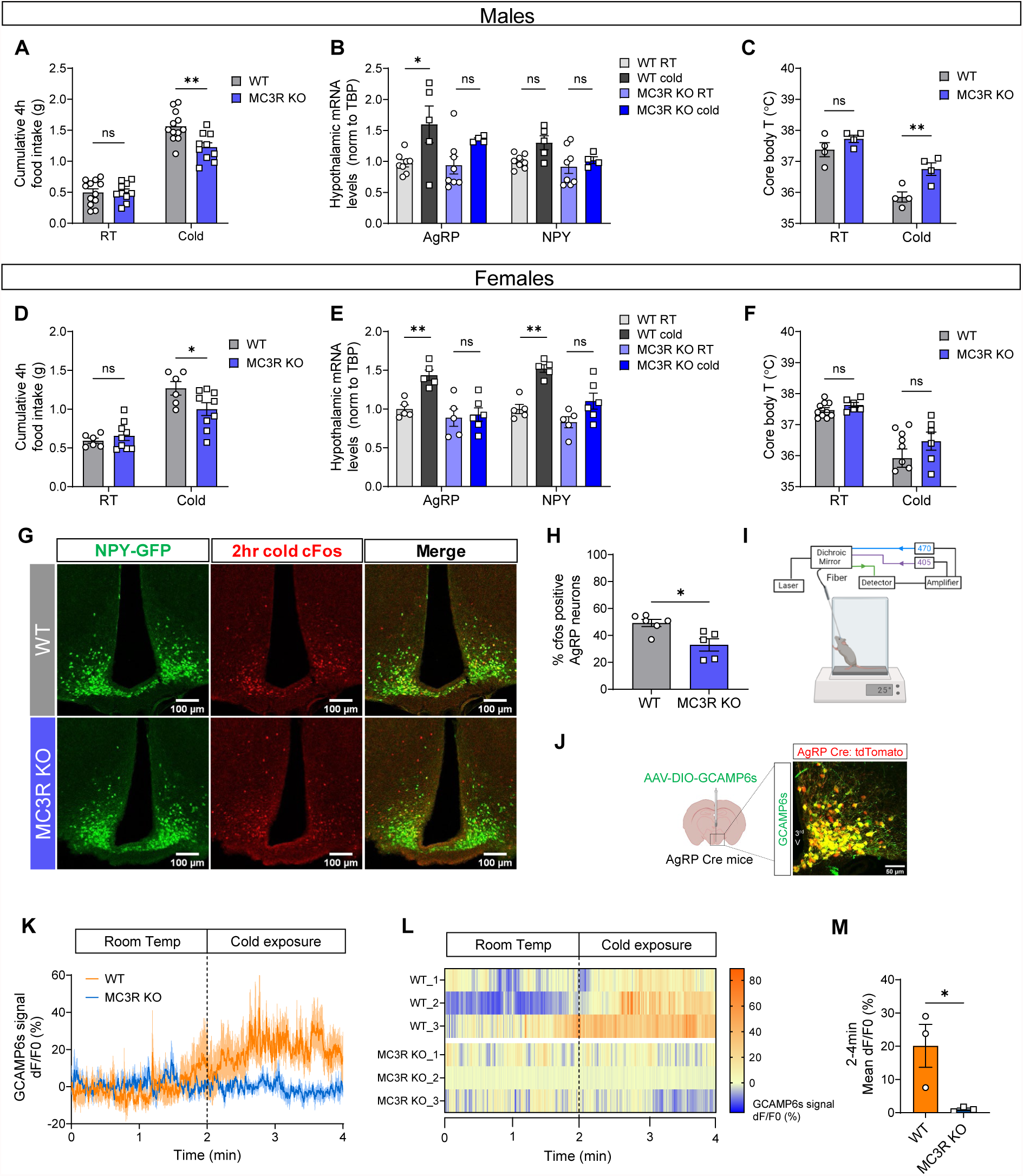
MC3R is required for the activation of AgRP neurons in response to cold. (A, D) Four-hour food intake of WT and MC3R KO male and female mice under room temperature (RT, 22°C) or cold exposure (4°C) (n = 6∼12 mice for all groups). (B, E, C, F) qPCR analysis of hypothalamic AGRP and NPY mRNA levels (B, E) and measurement of rectal temperature (C, F) in WT and MC3R KO male and female mice under room temp or following 4h cold exposure (n = 4∼10 mice for all groups). (G, H) Representative images of GFP and cFos immunostaining, and quantifications of the percentage of cFos-positive GFP cells in the ARC of WT and MC3R KO NPY-GFP male mice following 2h cold exposure. Scale bar, 100μm (n = 5∼6 mice for each group). (I) Schematic showing the setup of fiber photometry on a cold plate. (J) Validation of Cre-dependent GCAMP6s viral expression in AgRP Cre: tdTomato mice. Scale bar, 50μm. (K, L, M) Traces, heatmap and quantifications of averaged dF/F0 (%) GCaMP6s signal in AgRP neurons in WT and MC3R KO male mice. 0-2min, 22°C; 2-4min, 12°C (n=3 mice for each group). Data are plotted as mean and all error bars represent the SEM. ns, non-significant; *p < 0.05; **p < 0.01; ***p < 0.001, ****p < 0.0001 in unpaired Student’s t-test and two-way ANOVA with Sidak’s posthoc test.

We next performed cFos immunostaining and fiber photometry to directly examine AgRP neuron activity following cold exposure. Compared to WT mice that demonstrated evident cold-induced cFos expression in AgRP neurons, MC3R KO male mice showed significantly less cFos signal following cold exposure (Figures 2G and 2H). To measure calcium activity in AgRP neurons, freely moving mice that expressed GCaMP6s in AgRP neurons were placed onto a temperature-controlled metal plate (Figures 2I and 2J). In accordance with previous findings,^23, 24^ AgRP neuron activity in WT animals rapidly increased following the drop in environmental temperature (22 °C to 12°C). However, such cold-induced increase in AgRP neuron activity was greatly blunted in MC3R KO mice (Figures 2K-2M). Together, these findings suggest MC3R is required for the activation of AgRP neurons in sensing different forms of energy need, including not only food deprivation but also cold challenge.

### MC3R is not required for the inhibition of AgRP neurons by sensory detection of food cues

Having demonstrated the role of MC3R in regulating AgRP neuron activation by signals of energy need, we next asked if MC3R is also required for the inhibition of AgRP neurons in response to food cues or ingestion of nutrients, as shown previously.^25–28^ We investigated how MC3R deletion affects inhibition of AgRP neurons in hungry mice, in response to either caged or accessible chow. Importantly, the mRNA analysis from Figures 1E-1J revealed delayed AgRP mRNA upregulation in MC3R KO mice upon fasting, suggesting the importance of normalizing the extent to which AgRP neurons are activated in two groups before examining the inhibition of AgRP neurons by food cue. To this end, we paired 12hr-fasted WT to 36hr-fasted MC3R KO male mice that exhibited comparable AgRP mRNA levels in Figure 1E. We found that in response to “caged” or accessible food pellets, 36hr-fasted MC3R KO mice exhibited equal levels of decrease in AgRP neuron activity compared to 12hr-fasted WT group (Figures S2A-2D). These results suggested that despite compromised activation of AgRP neurons in the absence of MC3R, MC3R KO mice exhibit intact kinetics of AgRP neuron inhibition by sensory detection of food cues.

We next examined the requirement of MC3R in highly palatable food-induced inhibition of AgRP neurons in *ad lib* fed mice by performing oral gavage of Ensure. As previously described,^25, 28^ intragastric nutrient detection rapidly and durably inhibited AgRP neuron activity in a caloric-dependent manner. We found that in response to 30% or 100% Ensure gavage, WT and MC3R KO mice again exhibited comparable decrease in AgRP neuron activity (Figures S2E-2H). Taken together, our data suggested that MC3R is not required for the inhibition of AgRP neurons by sensory detection of food cues or nutrients.

### MC3R regulates the activation of AgRP neurons in a cell-autonomous manner

Given the dense expression of *MC3R* transcripts detected in AgRP neurons,^12^ we next asked if MC3R-mediated regulation of AgRP neurons is cell-autonomous. To this end, we first validated the expression of MC3R protein in AgRP cell bodies and AgRP nerve terminals in the paraventricular nucleus of hypothalamus (PVN). We unilaterally injected Cre-dependent virus coding a flag-tagged MC3R protein into the ARC of AgRP Cre mice. Immunohistochemical detection of FLAG expression colocalized with AgRP cell bodies in the ARC and with AgRP nerve terminals in the PVN by immunostaining (Figures 3A-C).

**Figure 3.**
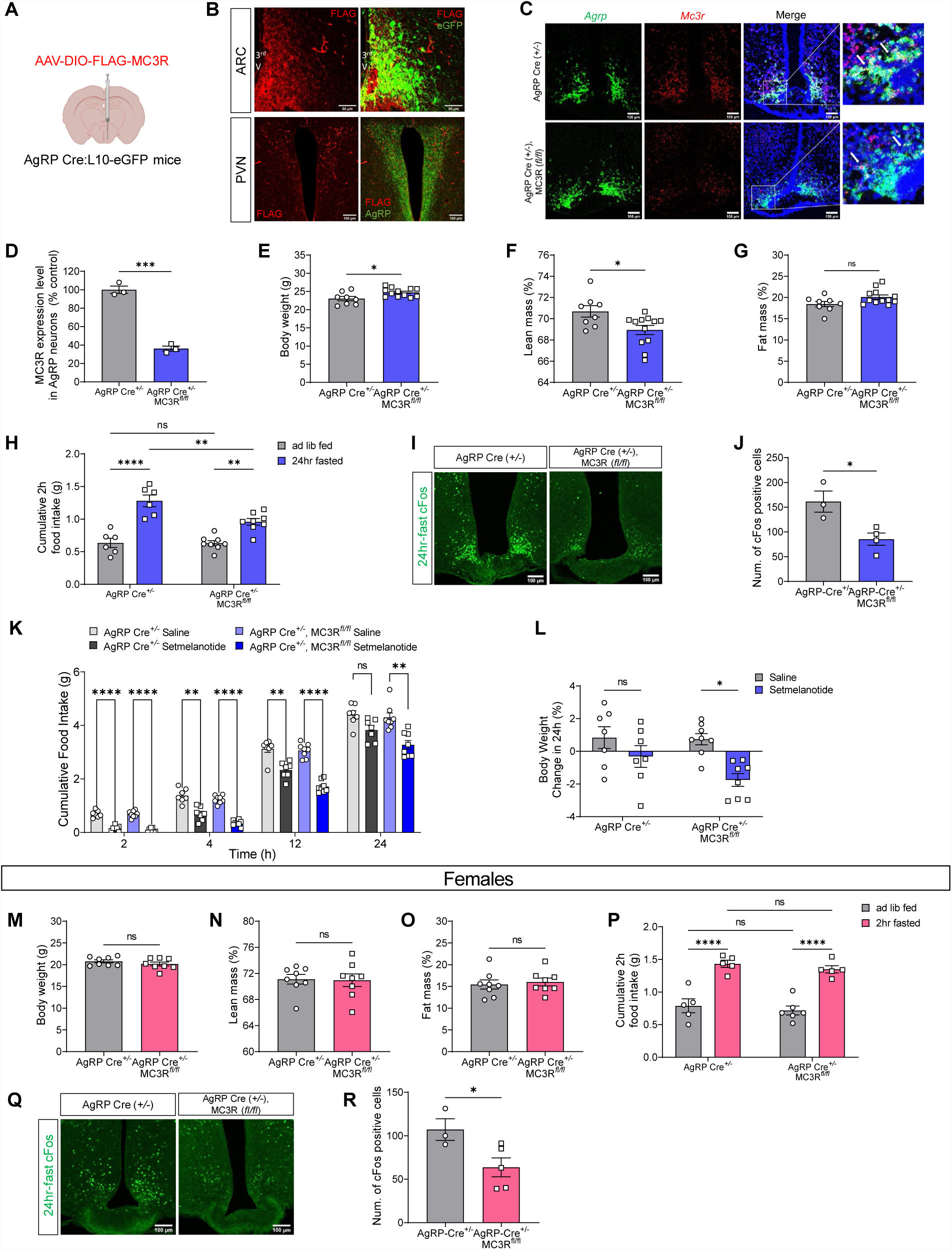
MC3R regulates the activation of AgRP neurons in a cell-autonomous manner. (A, B) Validation of Cre-dependent MC3R-FLAG viral expression in AgRP Cre: L10-eGFP mice by immunostaining against FLAG, eGFP and AgRP. Scale bar, 50μm for the ARC and 100μm for the PVN. (C, D) RNAscope analysis of *Mc3r* and *Agrp* mRNA expression, and quantifications of the percentage of AgRP-positive cells co-expressing *Mc3r* in the ARC of AgRP-Cre and AgRP-specific MC3R KO male mice. Scale bar, 100um (n = 3 mice for each group). (E-G, M-O) Body weight and body composition of AgRP-Cre and AgRP-specific MC3R KO male and female mice (n = 8∼12 mice for all groups). (H, P) Two-hour food intake of AgRP-Cre and AgRP-specific MC3R KO male and female mice following a 24hr fast (n=5∼8 mice for all groups). (I, J, Q, R) Representative images of cFos immunostaining, and quantifications of cFos-positive cell number in the ARC of 24h-fasted AgRP-Cre and AgRP-specific MC3R KO male and female mice. Scale bar, 100μm (n = 3∼5 mice for all groups). (K, L) Food intake and percent change 24h body weight in AgRP-Cre and AgRP-specific MC3R KO male mice given saline or setmelanotide injection (2mg/kg, i.p.) (n = 7∼8 mice for all groups). Data are plotted as mean and all error bars represent the SEM. ns, non-significant; *p < 0.05; **p < 0.01; ***p < 0.001, ****p < 0.0001 in unpaired Student’s t-test and two-way ANOVA with Sidak’s posthoc test.

Next, we sought to specifically delete MC3R in AgRP neurons by crossing AgRP-Cre with MC3R-flox mouse lines to generate a conditional MC3R KO model. Using RNAScope *in situ* hybridization, we validated knockout efficiency by showing *Mc3r* deletion in ∼70% of AgRP neurons (Figure 3D). We then interrogated whether the mild obesity and the defective fasting responses in global MC3R KO mice could be recapitulated by AgRP-specific deletion of MC3R. We found that AgRP-specific MC3R KO male mice exhibited increased body weight and decreased lean mass percent compared to the AgRP-Cre control group (Figure 3E-G, which phenocopied the global MC3R KO model.^6, 7^ In AgRP-specific MC3R KO females, however, there was no difference in their body composition compared to control group (Figure 3M-O). In response to fasting, we found that both male and female AgRP-specific MC3R KO mice showed defective cFos expression in the ARC (Figure 3, I-J, Q-R), while only males but not females demonstrated blunted refeeding behavior compared to control group (Figure 3, H, P).

Previous electrophysiological data supported that MC3R is required for GABA release from AgRP neuron terminals onto downstream PVN MC4R neurons,^11^ implying the role of MC3R as a negative regulator of the MC4R circuitry. In line with this, global MC3R KO mice exhibited hypersensitivity to MC4R agonists.^11^ We then sought to examine the effect of AgRP-specific MC3R deletion on MC4R-mediated anorexigenic responses. To this end, we chose setmelanotide, an FDA-approved MC4R agonist drug for treatment of POMC and leptin receptor deficiency, ^29^ and tested its anorexigenic action over a 24h time course following injection. Compared to the control group, the administration of setmelanotide (2mg/kg, i.p.) in AgRP-specific MC3R KO mice generated more robust and longer-lasting effect in food intake suppression at all time points measured after 1 hour (Figure 3K). Consistently, setmelanotide induced more profound weight loss in AgRP-specific MC3R KO group compared to control mice (Fig. 3L). Thus, our findings confirmed the hypersensitivity of downstream anorexigenic MC4R neurons in the absence of MC3R in AgRP neurons.

### Defective AgRP neuron activation in MC3R KO mice is not a result of developmental adaptation or differential reduction in leptin

We next investigated the mechanisms underlying the defective AgRP neuron activation in the absence of MC3R. One possibility is that such a defect is a result of developmental adaptions to MC3R loss. Although a previous study showed that the defective fasting-induced refeeding can be reproduced via knockdown of arcuate *Mc3r* expression in adult mice,^11^ the activation of AgRP neurons was not examined. To test this, here we performed viral-mediated knockdown of *Mc3r* by injecting AAV coding short hairpin RNA (shRNA) against *Mc3r* or a scrambled control sequence unilaterally in the ARC of adult WT male mice (Figure S3A). Following shRNA expression, we performed cFos immunostaining to compare the number of fasting-activated neurons in the viral injection side versus the control side (Figure S3B). In scramble shRNA-expressing mice, cFos signals were equally expressed in the two sides regardless of fed or fasted state. In *MC3R* shRNA-expressing mice, however, fasting-induced cFos expression was significantly less in the viral injection side compared to the control side (Figures S3C and S3D). These results suggested that the impaired AgRP neuron activation in MC3R KO mice could be recapitulated by ARC *Mc3r* knockdown in adulthood. Therefore, the defective fasting-induced activation of AgRP neurons is not a result of developmental adaptations to MC3R loss. Notably, despite normal body weight compared to scramble shRNA-expressing mice, *Mc3r* shRNA-expressing mice exhibited decreased fat mass percent and increased lean mass percent (Figures S3E-S3G), consistent with a role of arcuate MC3R as a negative regulator of the anorexigenic melanocortin circuits.

Another possibility that could explain the defective AgRP neuron activation in the absence of MC3R is that MC3R KO mice have a defect in fat utilization. In response to fasting, adipose tissues undergo lipolysis to provide sources as energy fuel, accompanied by decrease in plasma leptin levels.^30^ Previous work has shown that such a decrement in leptin levels is critical for AgRP neuron activation during fasting.^31^ To test for a defective fast-induced mobilization of adipose mass, we measured fasting-induced changes in fat mass and leptin levels in MC3R KO and WT male mice. Despite hyperleptinemia, we found that MC3R KO mice exhibited a comparable decrease in plasma leptin levels in response to a 24h fast compared to WT group (Figures S3H, S3I and S3L). In addition, the linear regression analysis of decreased fat mass and decreased leptin levels during fasting demonstrated comparable results between MC3R KO and WT groups (Figures S3J and S3K), further suggesting that MC3R KO mice have normal fasting-induced fat utilization. In AgRP-specific MC3R KO mice, we also observed equal plasma leptin levels and comparable linear relationship between leptin levels and fat mass compared to control group (Figures S3M and S3N). Taken together, our data suggested that defective AgRP neuron activation in MC3R KO mice is not a result of differential reduction in leptin.

### MC3R is required for the orexigenic action of ghrelin in AgRP neurons

In addition to leptin, ghrelin is another hormonal signal of energy state that regulates AgRP neuron activity.^32^ Inadequate sensing of ghrelin could be another mechanism underlying defective AgRP neuron activation in MC3R KO mice. Previous work has shown that compared to WT mice, global MC3R KO mice exhibit normal serum ghrelin levels but eat significantly less in response to ghrelin injection.^33^ Here, we first sought to confirm the requirement of AgRP-specific MC3R expression for such ghrelin-induced hyperphagia. To this end, we examined food intake following peripheral ghrelin administration in the presence and absence of MC3R in AgRP neurons. In the control group, ghrelin injection (1.6mg/kg, i.p.) significantly increased acute food intake. In AgRP-specific MC3R KO mice, however, ghrelin-induced hyperphagia was diminished (Figure 4A), confirming that MC3R is required for the orexigenic effect of ghrelin through its cell-autonomous action in AgRP neurons.

**Figure 4.**
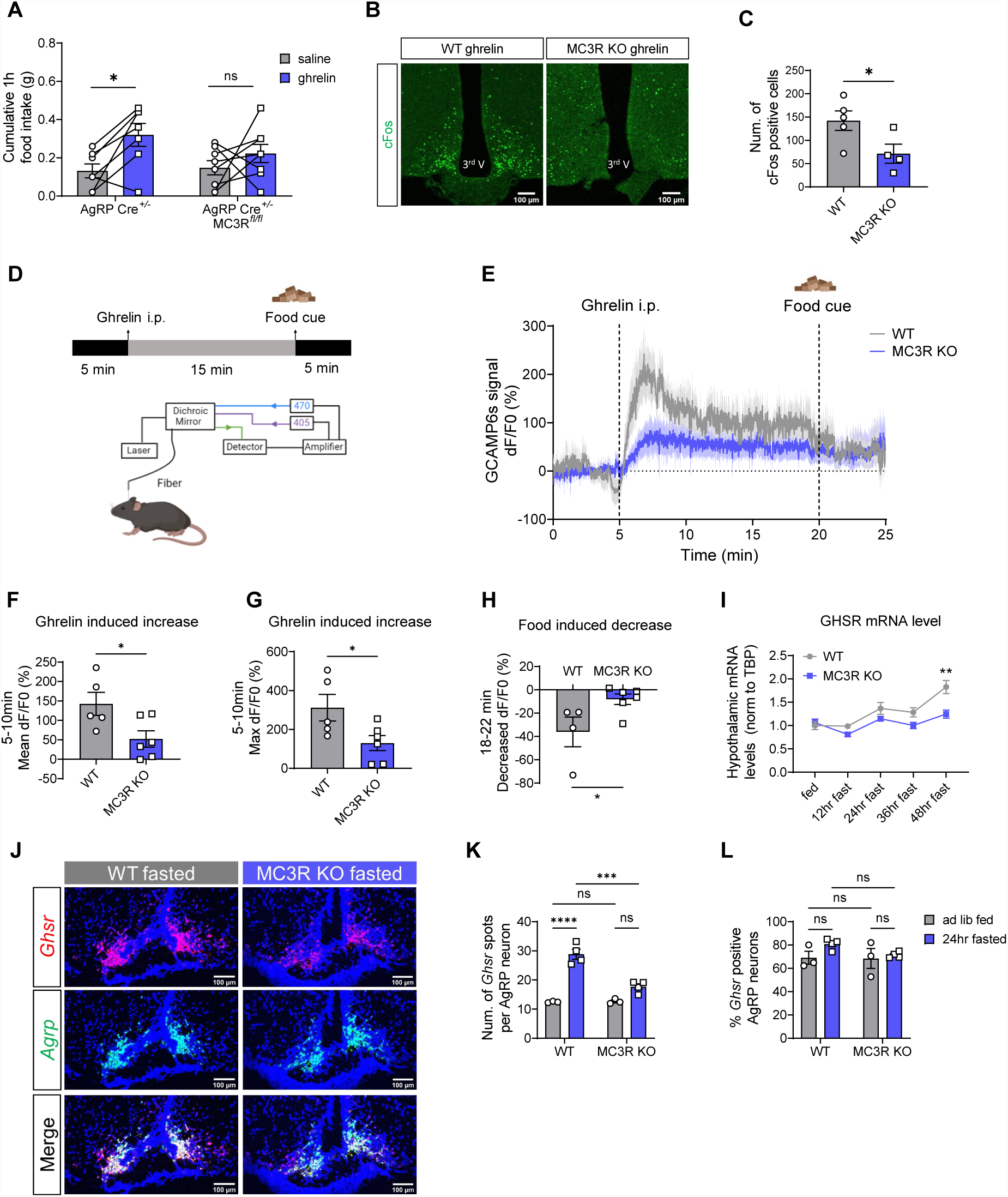
MC3R is required for the orexigenic action of ghrelin on AgRP neurons. (A) One-hour food intake of AgRP-Cre and AgRP-specific MC3R KO male mice given saline or ghrelin injection (1.6 mg/kg, i.p.) (n = 7∼8 mice for all groups). (B, C) Representative images of cFos immunostaining, and quantifications of cFos-positive cell number in the ARC of WT of MC3R KO male mice in response to saline or ghrelin injection. Scale bar, 100um (n = 4∼5 mice for each group). (D) Schematic showing the time course of ghrelin injection and food presence in fiber photometry experiment. (E-H) Traces and quantifications of averaged dF/F0 (%) GCaMP6s signal in AgRP neurons in WT and MC3R KO male mice in response to ghrelin and food cue (n = 5∼6 mice for each group). (I) qPCR analysis of hypothalamic GHSR mRNA levels in WT and MC3R KO female mice over a 48h fasting time course (n = 3∼5 mice for all groups). (J-L) RNAscope analysis of *Agrp* and *Ghsr* mRNA expression, and quantifications of the number of *Ghsr* transcripts in each AgRP neuron and the percentage of AgRP neurons co-expressing *Ghsr* in the ARC of fed or 24h-fasted WT and MC3R KO male mice. Scale bar, 100um (n = 3∼4 mice for all groups). Data are plotted as mean and all error bars represent the SEM. ns, non-significant; *p < 0.05; **p < 0.01; ***p < 0.001, ****p < 0.0001 in unpaired Student’s t-test and two-way ANOVA with Sidak’s posthoc test.

We next examined the activation of AgRP neurons by ghrelin in the presence and absence of MC3R. We performed cFos immunostaining and found that ghrelin-induced cFos expression in the ARC was significantly suppressed in MC3R KO mice compared to WT group (Figures 4B and 4C). We then performed fiber photometry to record calcium signals in AgRP neurons in response to ghrelin. In WT mice, as previously described, ghrelin administration (1.6mg/kg, i.p.) induced a rapid and robust increase in AgRP neuron activity, which was greatly suppressed by subsequent sensory detection of a chow pellet (Figures 4D-H).^26^ In MC3R KO mice, however, we found a significant impairment in ghrelin-induced increase in AgRP neuron activity (Figure 4E-G) and accordingly less inhibition by subsequent food presence (Figure 4E, H).

We next investigated the effect of *Mc3r* deletion on ghrelin receptor expression under *ad lib* fed and fasted states. We performed RNAScope *in situ* hybridization to examine the levels of *Ghsr* mRNA in AgRP neurons in WT and MC3R KO male mice (Figure 4J-L). In the WT group, fasting significantly increased the number of *Ghsr* transcripts in AgRP neurons, while the percent of AgRP neurons expressing *Ghsr* was unchanged by fasting. In MC3R KO group, however, there was an evident defect in fasting-induced *Ghsr* upregulation in AgRP neurons. We also performed qPCR analysis of *Ghsr* mRNA levels and found that compared to WT group, MC3R KO mice exhibited significantly less upregulation of *Ghsr* expression in hypothalamus upon fasting (Figure I). Together, these findings suggested that MC3R deletion from AgRP neurons leads to impaired response to ghrelin, concomitant with a hypothalamic defect in fasting-induced upregulation of *Ghsr* mRNA levels.

## DISCUSSION

In this study, we demonstrate a cell-autonomous role for MC3R in regulating AgRP neuron activation in response to two different signals of energy demand: food deprivation and cold exposure. The data here show a clear requirement of MC3R expression within AgRP neurons for the normal activation of these cells in response to nutritional deficit. This finding, in turn, helps explain the multiple observations of defective fast-induced behavioral and neuroendocrine responses to fasting reported in the MC3R KO mouse.^11, 14–16^ We found that despite the requirement of MC3R for AgRP neuron activation, MC3R is dispensable for the inhibition of AgRP neurons, likely understood in terms of the different mechanism(s) by which AgRP neurons are activated versus deactivated.^25^

While the cellular basis for the effects of MC3R on the fasting response is described here, involving MC3R expression within AgRP neurons, the molecular basis remains unknown, and could involve either presynaptic, and/or or postsynaptic mechanisms of MC3R action. The presynaptic actions of the MC3R have been demonstrated, with slice electrophysiological studies showing a role for the MC3R in stimulating GABA release from AgRP termini onto both downstream melanocortin receptor expressing cell types tested, the MC3R-expressing POMC neurons,^13^ and the PVN MC4R neurons.^11^ The data presented here shows that AgRP neurons are still activated by fasting in the absence of MC3R function, but exhibit greatly reduced sensitivity to signals of nutritional deficit, such as a decrease in serum leptin, or an increase in ghrelin. Leptin levels are sensed not only by AgRP neurons, but also a through a network of LepRb-expressing GABAergic neurons^34^ such as the GABAergic DMH^LepRb^ neurons known to densely innervate AgRP neurons.^35^. The results shown here are consistent, for example, with increased GABAergic tone on the AgRP neurons, which could result if there are reciprocal connections between AgRP neurons, and other LepRb-expressing GABAergic neurons, with reduced GABAergic drive from MC3R-deficient AgRP termini.

While we show here that a tagged MC3R is expressed in AgRP soma, there is little other data to support a postsynaptic role for the MC3R on AgRP soma. Innervation of AgRP cell bodies by POMC or AgRP fibers has not been reported, and no responses were observed in AgRP cell bodies following optogenetic stimulation of POMC or AgRP neurons.^36^ Further, direct application of a broad melanocortin agonist, MTII, to AgRP cells was not seen to alter the activity of these neurons, although the endogenous antagonist of the MC3R and MC4R, AgRP, was not tested. ^37^ Thus, additional work is required to determine if endogenous MC3R is actually expressed on AgRP soma, and if so, what the source of the ligand is.

Remarkably, the deletion of a number of different receptors and receptor accessory proteins from AgRP neurons, including the NMDA receptor,^38^ growth hormone receptor,^39^ and GPCR accessory protein MRAP2^40^ all blunt the responsiveness of AgRP neurons to fasting-induced activation. The observation that deletion of any one of several different receptors or receptor accessory proteins from AgRP neurons (MC3R, GHR, MRAP2, NMDAR) each can blunt fasting-induced activation may suggest that these molecules are part of a large signaling complex, and perhaps the MC3R is required for the trafficking or function of the complex.

However, the results shown here for the MC3R are clearly different from these other receptors in that they further implicate the MC3R as a negative regulator of the control of AgRP neuron activity. Further, these findings make the MC3R a novel potential drug target for blocking weight regain, since MC3R-specific antagonists could be expected to blunt the behavioral and neuroendocrine responses to weight loss, regardless of whether such loss results from pharmacotherapy, or diet and exercise.

### Limitations of the study

In this study, we have used global and tissue specific deletion of the MC3R to characterize functions of this receptor, and thus it is possible that some phenotypes result from developmental activities of the receptor. However, we control for this by showing that knockdown of the ARC MC3R in adult mice also blunts fasting-induced activation of cFos in the ARC (Figure S3). While expression of MC3R protein in MC3R soma in the ARC can be shown by viral overexpression, this does not prove the endogenous MC3R is functionally expressed on AgRP soma. Ultimately, slice electrophysiology of AgRP neurons will be needed to test the hypothesis that the loss of MC3R in AgRP neurons alters GABAergic tone on these cells, or that endogenous MC3R is functionally expressed on MC3R soma.

## METHODS

### Animals

Experiments were performed on adult (12-16 weeks old) male and female mice. In this study, mouse strains including C57/BL6J (Jax: 000664), AgRP-IRES-Cre (Jax: 012899), NPY-hrGFP (Jax: 006417), Ai14 (Jax: 007914) and EGFP-L10a (Jax: 024750) were obtained from Jackson Laboratories. age. MC3R KO mice were generated and maintained in a C57BL/6 background as previously described ^7^. MC3R ^flox/flox^ (MC3R ^fl/fl^) mice are reported here for the first time. Briefly, MC3R ^fl/fl^ mice were made by inserting loxP sites to flank the single exon using a guide RNA (gRNA) to the Mc3R gene, with the donor vector containing the loxP sites (Cyagen, Inc). The 5′ loxP site is inserted into 5′ UTR and the 3′ loxP site sits after the 3′ UTR. The gRNA (target sequence: AAGAGTTCATGGTTAACAGCAGG) was co-injected with Cas9 to generate conditional knockout offspring. Founder animals were then bred to wild type to test for germline transmission. F1 heterozygous mutants were identified by additional PCR screening using two sets of primers: 5’arm forward primer (F2): 5’-GTTCATCTGTCTAGCAGCTTCATT-3’ 3’loxP reverse primer (R2): 5’-GTGGATTCGGACCAGTCTGA-3, and 5’loxP forward primer (F1): 5’- ACGTAAACGGCCACAAGTTC-3’ 3’arm reverse primer (R1): 5’- TTATCCACCACACCTTGCTTTCTC-3’. For sequencing confirmation, additional PCR was conducted using 2 sets of primers: Sequencing Primer for PCR product 1: 5’Sequence primer (R4): 5’-GATAAACTGGTCCTCCAAGGTCAG-3’ Sequencing Primer for PCR product 2: 3’Sequence primer (R3): 5’-GGTAGAATCTCCTTGTGTGTGTTT-3’. MC3R ^fl/fl^ mice were bred to AgRP-Cre to produce AgRP-specific MC3R KO mice (AgRP Cre ^+/-^, MC3R ^fl/fl^). All experiments were previously approved by the University of Michigan Institutional animal care and use office (IACUC).

### Food intake studies

For all feeding and behavioral assays mice were matched for gender, age, and body weight. Gender differences in behavioral assays are depicted in either the main text or the supplementary figures. Mice were provided with *ad libitum* access to food and water in temperature-controlled (22°C) rooms on a 12 h light-dark cycle with daily health status checks. For feeding assays mice were single housed for at least two weeks before starting feeding measurements. Food pellets were provided in a wire feeder, and intake was measured manually including all crumbs.

### Fast-refeeding assays

12 to 16-week-old WT and MC3R KO male and female mice were fasting for 24 hours. Right before the dark cycle, a weighed amount of food was provided, and the food was weighed again 2hr, 4hr, 8hr and 24hr after refeeding. Food intake was calculated as the difference in food weight over the 24hr refeeding time course, and was compared to the *ad lib* fed food intake in the same time period.

### Cold-exposure feeding assays

Single-cage housed WT and MC3R KO mice were habituated to the temperature-controlled chamber at room temperature (22 °C) for 3 days prior to the test. For the cold-induced feeding experiment, mice were maintained in the chamber at 4 or 22 °C with free access to food and water for 4hr in the light cycle. The food was weighted right before and after the cold exposure.

### Real-Time PCR Analyses

To measure the gene expression of *Agrp*, *Npy*, *Pomc* and *Ghsr*, WT or MC3R KO mice were *ad lib* fed or fasted for 12, 24, 36 and 48 hours from the beginning of the dark cycle. To measure the gene expression in response to cold exposure, acclimated mice were maintained in the chamber at 4 or 22 °C with free access to water but food deprived for 4hr in the light cycle. Following fasting or cold exposure, mice were killed and hypothalamic tissue were collected and frozen immediately on dry ice and stored at −80 °C. RNA was extracted using TRIzol reagent (Invitrogen), followed by cDNA synthesis using the cDNA reverse transcription kit (Applied Biosystems) according to the manufacturer’s instructions. SYBR Green assay was used to examine the gene-expression levels of *Agrp*, *Npy*, *Pomc*. Gene expression for each target was normalized to TATA box binding (*Tbp*) gene.

### Immunofluorescence and Imaging

To measure changes in AgRP neuronal activity in response to fasting or cold exposure, we performed florescent immunohistochemistry to detect cFos and GFP. Mice were euthanized and perfused with 1X PBS followed by 10% formalin. Following perfusion, brains were fixed for an additional 24 hours in 10% formalin. Brains were then switched to a 30% sucrose solution (in 1X PBS) until the brains sank in the solution (1-3 days) and at which point 35um thick hypothalamic sections were obtained using a cryostat (Leica). Sections were first incubated for 1 hour in blocking buffer (1X PBS with 2% BSA and 0.1% tween-20). Primary antibodies (rabbit anti-cFos, Cell Signaling Technology, 1:500; chicken anti-GFP, Aves Labs, 1:1500) diluted in blocking buffer were added to sections overnight at room temp. After three washes in 1X PBS, secondary antibodies (goat anti-chicken Alexa 488, anti-rabbit Alexa 555, 1:500) diluted in blocking buffer at 1:500 were added to sections for 2 hours at room. After three additional wash steps, sections were mounted on slides and imaged on a confocal microscope (Leica SP8). Total number of cFos+ cells, GFP+ cells, cFos+/GFP+ dual positive cells in the ARC were counted. The percentage of cFos-positive AgRP neurons and GFP-positive cFos cells were calculated accordingly.

For MC3R expression validation experiment, FLAG and AgRP primary antibodies were used and diluted in blocking buffer at 1:200 (mouse anti-FLAG, Sigma) and 1:1000 (rabbit anti-AgRP, Phoenix Pharmaceuticals), respectively. Secondary antibodies (goat anti-mouse Alex 555, anti-rabbit Alexa 647) were diluted in blocking buffer at 1:500 accordingly.

### Rectal thermometry

A small-diameter temperature probe was inserted through the anus in WT and MC3R KO mice, with an insertion depth of >2 cm to yield colonic temperatures right before and after cold exposure.

### Viral vectors

The adeno associated viral vector AAV5-synapsin-DIO-GCAMP6s used in this study was purchased from Addgene. The AAV9-CMV-DIO-FLAG-MC3R, AAV2-CMV-MC3R shRNA and AAV2-CMV-scramble shRNA were designed by the University of Michigan Vector Core. The generation and the sequences of the MC3R and scramble shRNAs have been described previously ^10^.

### Stereotaxic viral injections and fiber placement surgeries

For stereotaxic surgical procedures, 8 to 12-week-old mice were anesthetized with isoflurane and placed in a stereotaxic frame (Kopf). A micro-precision drill was used to drill a small burr-hole directly above the viral injection point and dura was removed. AAV viral vectors of 500nl were injected into the arcuate nucleus using a micromanipulator (Narishige) attached to a pulled glass pipette. Viral injection coordinates for the arcuate nucleus were as follows: A/P: −1.4mm (from bregma), M/L: 0.25mm, D/V: 5.95mm (from surface of the skull). Virus was injected at a rate of 50nl/minute. Following viral injection, the glass pipette was left in place for an additional ten minutes to prevent leaking of virus outside the targeted brain region. For fiber implant surgery, after viral injections a fiber-optic ferrule was implanted using the coordinates: A/P: −1.4mm (from bregma), M/L: 0.25mm, D/V: 5.85mm (from surface of the skull). Fibers were attached to the skull using Metabond. Three weeks was allotted post-surgery to allow for viral expression and the mice to recover from surgical procedures. Viral expression and fiber placement were verified post hoc in all animals, and any data from animals in which the transgene expression and/or fiber was located outside the targeted area were excluded from analysis.

### Fiber photometry

Mice expressing GCaMP6s in AgRP neurons were connected to a fiber-photometry system to enable fluorometric analysis of real-time neuronal activity. Data were recorded using a Neurophotometrics FP3001 system (Neurophotometrics, San Diego, CA). Briefly, 470 and 415 nm LED light was bandpass filtered, reflected by a dichroic mirror, and focused onto a multi-branch patch cord (Doric, Quebec City, Quebec) by a 20x objective lens. Alternating pulses of 470 and 415 nm light (∼50 µW) were delivered at 40 Hz, and photometry signals were analyzed using custom MATLAB scripts. For all recordings, the isosbestic 415 nm excitation control signal was subtracted from the 470 nm excitation signal to remove movement artifacts from intracellular calcium-dependent GCaMP6s fluorescence. Baseline drift was evident in the signal due to slow photobleaching artifacts, particularly during the first several minutes of each recording session. A double exponential curve was fit to the raw trace of temperature-ramping experiments while a linear fit was applied to the raw trace of food presentation experiments and subtracted to correct for baseline drift. After baseline correction, dF/F0 (%) was calculated as individual fluorescence intensity measurements relative to mean fluorescence of the beginning control session for 470 nm channel. In cold exposure experiments, the beginning control session was the 0-2min baseline recording at room temperature. In food detection and ghrelin response experiments, the beginning control session was the 0-5min baseline recording before any stimulations were presented.

### RNAscope fluorescent *in situ* hybridization

To determine the percentage of AgRP neurons that contain *Mc3r* or *Ghsr* and to quantify *Mc3r* or *Ghsr* mRNA abundance in individual AgRP neurons in the ARC, an RNAscope fluorescent multiplex assay (Advanced Cell Diagnostics) was used. Fresh brain samples were collected, embedded in OCT and frozen immediately on dry ice and stored at −80 °C. 18um-thick frozen sections containing the ARC were collected using a sliding microtome and mounted onto Super Frost Plus slides (Fisher Scientific), and *in situ* hybridization was performed according to the RNAscope fluorescent multiplex kit user manual for fresh frozen tissue (Advanced Cell Diagnostics), using RNAscope Probe-Mm-Mc3r-C1(Cat# 412541) or RNAscope Probe-Mm-Ghsr-C1(Cat# 426141), and RNAscope Probe-Mm-Agrp-C2 (Cat# 400711-C2). Images of the ARC of each animal were obtained using a laser scanning confocal microscope (Leica SP8). Confocal image stacks were collected through the z-axis. Total number of cells (boundary defined by DAPI signals) that co-express *Agrp* and *Mc3r* or *Ghsr*, and the number of *Mc3r* or *Ghsr* transcripts in each AgRP-positive cell were counted manually aided by ImageJ software. Only cells with three times as many fluorescent spots as were measured in background regions, were considered positively labeled for *Agrp*, *Mc3r* or *Ghsr* mRNA.

### Body composition

Whole-animal body composition analysis was performed using the Minispec Model mq7.5 (7.5 mHz; Bruker Instruments), which is a benchtop-pulsed 7-T NMR system. The instrument gave lean mass, fat mass, and fluid mass values of the mice, of which the body weight was measured at the same time. The percent of lean or fat mass relative to the whole-body weight was calculated.

### Plasma leptin measurements

Plasma leptin levels were measured in 24h-fasted and fed mice with enzyme-linked immunosorbent assay (Mouse Leptin ELISA Kit, Crystal Chem). Four to five drops of blood were collected in heparinized capillary blood collection tubes (Thermo Fisher Scientific) by tail vein puncture. Blood cells were removed by centrifugation (2000g for 30min at 4°C), and plasma was flash-frozen for further analysis according to the manufacturer’s recommendations.

### Setmelanotide injection

Male mice of each genotype (AgRP-speific MC3R KO or AgRP Cre) were weighed and divided into two subgroups, and each subgroup was given either vehicle (200 μl of saline) or setmelanotide (2 mg/kg) in 200 μl of saline by i.p. injection half an hour before the beginning of the dark phase. A weighed amount of food was provided, and the food was weighed again 2hr, 4hr, 12hr and 24hr after the injection. Mice were weighed again 24hr after injection to calculate the percent change in body weight.

### Ghrelin injection

Male mice of each genotype (AgRP-speific MC3R KO or AgRP Cre) were weighed and divided into two subgroups, and each subgroup was given either vehicle (200 μl of saline) or ghrelin (1.6 mg/kg) in 200 μl of saline by i.p. injection at 11am during the light phase (7am to 7pm). A weighed amount of food was provided, and the food was weighed again 1hr after the injection.

To test ghrelin-induced cFos expression, ghrelin was given by i.p. injection to *ad lib* fed mice at 11am during the light phase (7am to 7pm). The food was removed after ghrelin injection while mice had normal access to the water. Two hours following the injection, mice were euthanized and perfused with 1X PBS followed by 10% formalin for further cFos immunostaining experiment as described in the “Immunofluorescence and imaging” section.

## SUPPLEMENTAL INFORMATION

**Figure S1. MC3R is required for the activation of AgRP neurons in hunger sensing.**

**Figure S2. MC3R is not required for the inhibition of AgRP neurons by sensory detection of food cues or nutrients.**

**Figure S3. Defective AgRP neuron activation in MC3R KO mice is not a result of developmental adaptations or differential reduction in leptin.**

## ACKNOWLEDGEMENTS

We would like to thank Savannah Y. Williams and Monica Blazevic for their outstanding technical assistance. This work was funded by NIH DK126715 (R.D.C.), T32 DK101357 (N.S.D.), and K99DK127065 (P.S.).

## AUTHOR CONTRIBUTIONS

Conceptualization, R.D.C., P.S., and Y.G.; methodology, R.D.C., P.S., and Y.G.; investigation, Y.G., N.S.D., and G.D.; writing, Y.G. and R.D.C.; funding acquisition, R.D.C., P.S., and N.S.D.; supervision, R.C. and P.S..

