## Supplemental Figures for "Cell autonomous regulation of the activation of AgRP neurons by the melanocortin-3 receptor"

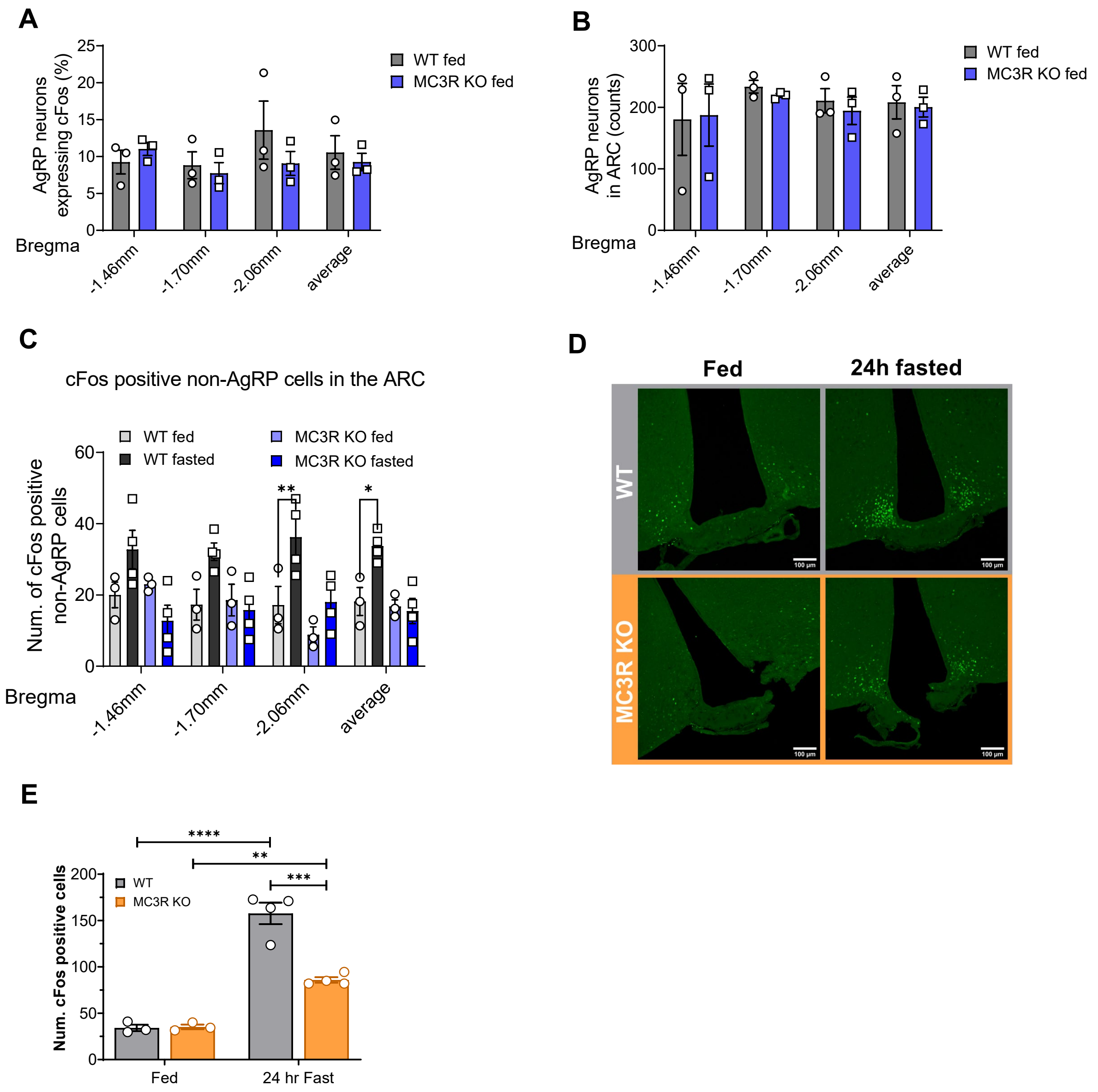


**Figure S1. MC3R is required for the activation of AgRP neurons in hunger sensing.**

(A, B) Quantifications of the percentage of cFos-positive and the number of GFP cells in the ARC of *ad lib* fed WT and MC3R KO NPY-GFP male mice (n = 3 mice for all groups).

(C) Quantifications of the number of cFos positive non-GFP cells in the ARC of WT and MC3R KO NPY-GFP, fed or 24h-fasted male mice (n=3~4 mice for all groups).

(D) Representative images of cFos immunostaining in WT and MC3R KO fed or 24h-fasted female mice. Scale bar, 100 um.

(E) Quantifications of the number of cFos-positive cells in the ARC of WT and MC3R KO fed or 24h-fasted female mice (n = 3~4 mice for all groups).

Data are plotted as mean and all error bars represent the SEM. ns, non-significant; $*p < 0.05$; $**p < 0.01$; $***p < 0.001$, $****p < 0.0001$ in two-way ANOVA with Sidak's posthoc test.

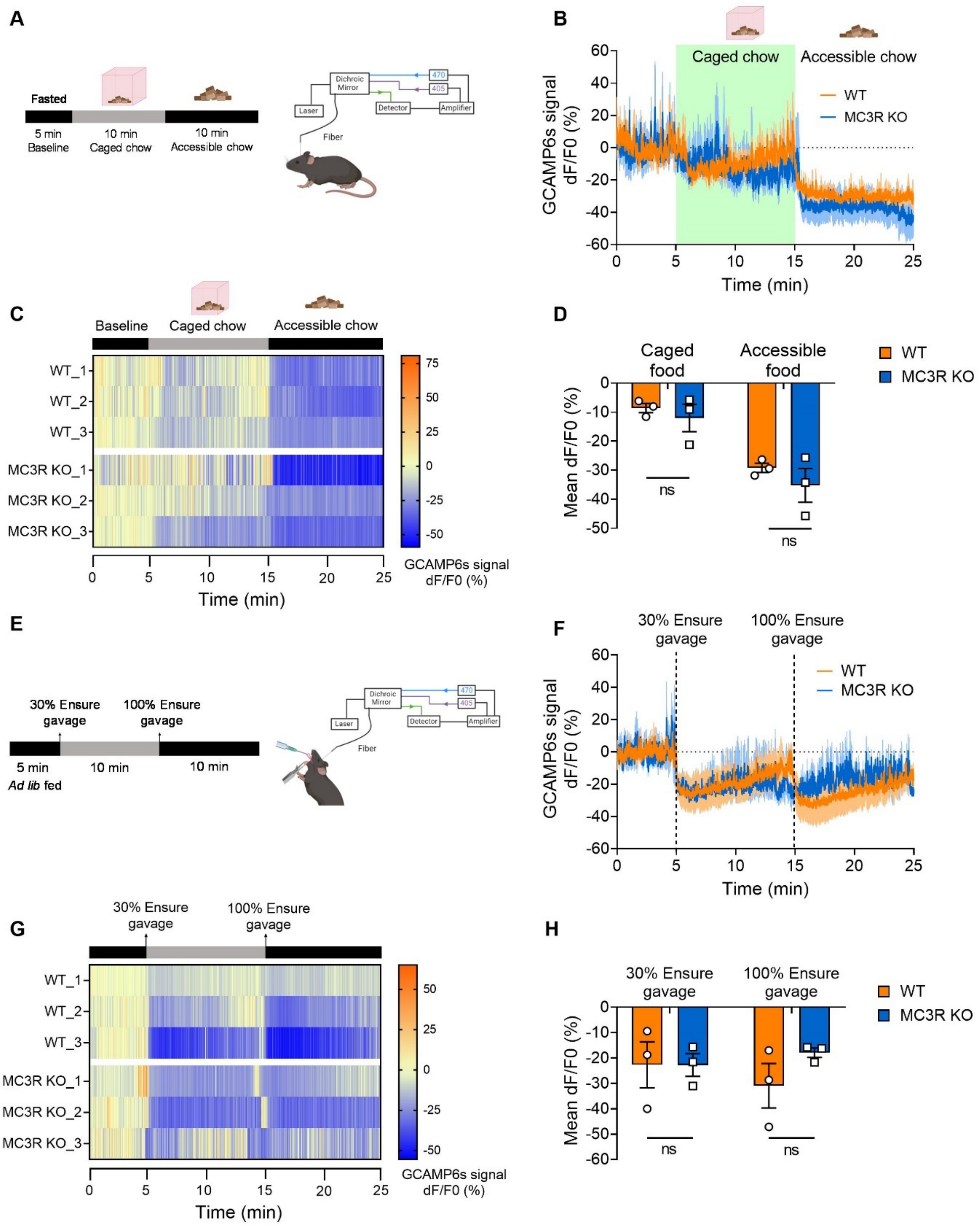
A
Fasted
5 min
Baseline
10 min
Caged chow
10 min
Accessible chow
Dichroic Mirror
470
405
Laser
Detector
Amplifier
Fiber
B
Caged chow
Accessible chow
WT
MC3R KO
GCAMP6s signal dF/F0 (%)
Time (min)
C
Baseline
Caged chow
Accessible chow
WT_1
WT_2
WT_3
MC3R KO_1
MC3R KO_2
MC3R KO_3
GCAMP6s signal dF/F0 (%)
Time (min)
D
Caged food
Accessible food
WT
MC3R KO
Mean dF/F0 (%)
ns
ns
E
30% Ensure gavage
100% Ensure gavage
5 min
Ad lib fed
10 min
10 min
Dichroic Mirror
470
405
Laser
Detector
Amplifier
Fiber
F
30% Ensure gavage
100% Ensure gavage
WT
MC3R KO
GCAMP6s signal dF/F0 (%)
Time (min)
G
30% Ensure gavage
100% Ensure gavage
WT_1
WT_2
WT_3
MC3R KO_1
MC3R KO_2
MC3R KO_3
GCAMP6s signal dF/F0 (%)
Time (min)
H
30% Ensure gavage
100% Ensure gavage
WT
MC3R KO
Mean dF/F0 (%)
ns
ns

**Figure S2. MC3R is not required for the inhibition of AgRP neurons by sensory detection of food cues or nutrients.**

(A) Schematic showing the setup of fiber photometry in fasted mice in response to caged or accessible food stimulation.

(B, C) Traces and heatmap of averaged dF/F0 (%) GCaMP6s signal in AgRP neurons in 12h-fasted WT or 36h-fasted MC3R KO male mice in response to food cues.

(D) Quantifications of decreased mean dF/F0 (%) GCAMP6s signal in response to caged or accessible food cue ($n = 3$ mice for all groups).

(E) Schematic showing the setup of fiber photometry in *ad lib* fed mice in response to 30% or 100% Ensure gavage.

(F, G) Traces and heatmap of averaged dF/F0 (%) GCaMP6s signal in AgRP neurons in *ad lib* fed WT or MC3R KO male mice in response to Ensure gavage.

(H) Quantifications of decreased mean dF/F0 (%) GCAMP6s signal in response to 30% or 100% Ensure gavage ($n = 3$ mice for all groups).

Data are plotted as mean and all error bars represent the SEM. ns, non-significant; *$p < 0.05$; **$p < 0.01$; ***$p < 0.001$, ****$p < 0.0001$ in two-way ANOVA with Sidak's posthoc test.

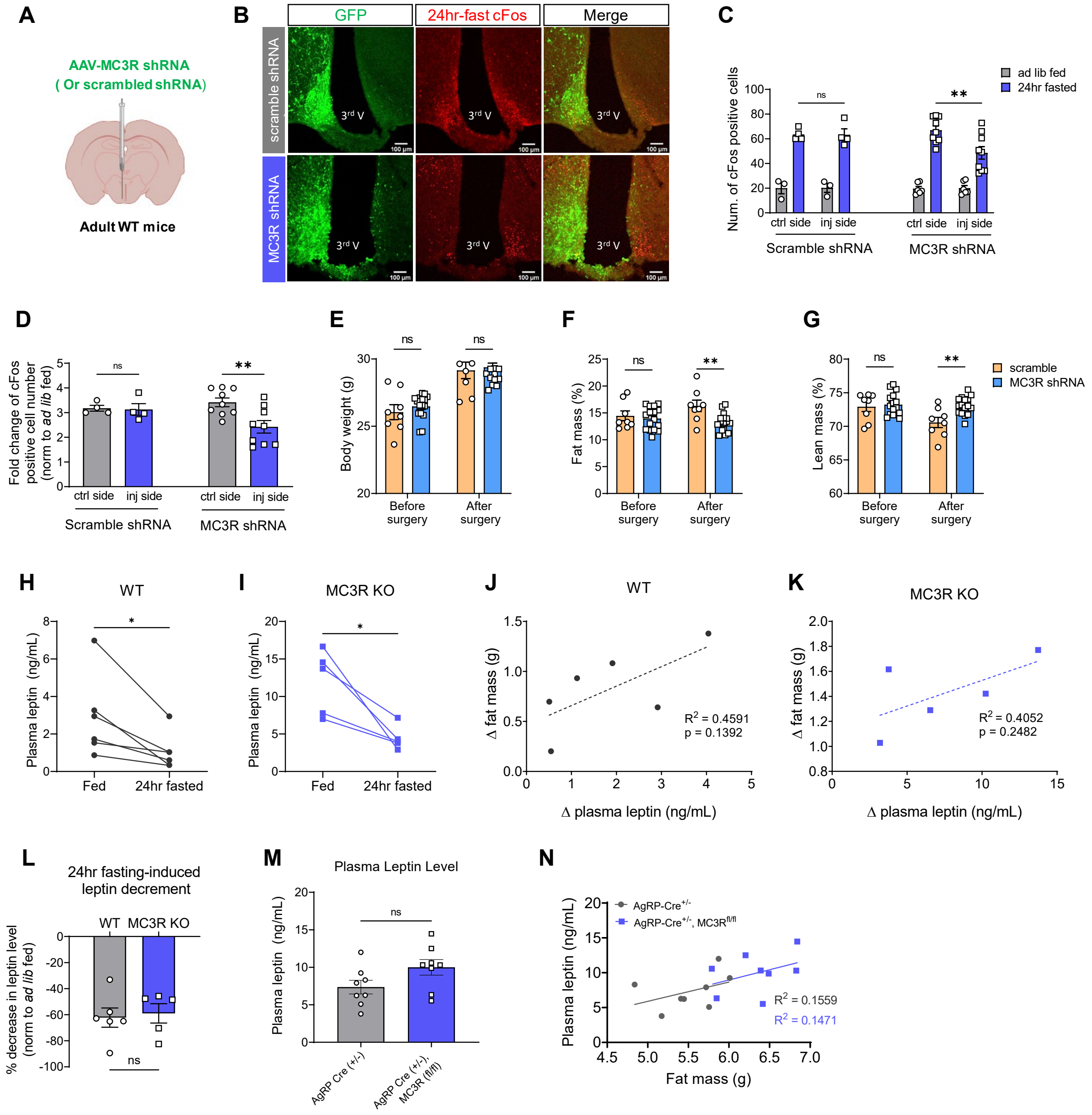

A
AAV-MC3R shRNA
( Or scrambled shRNA)
Adult WT mice
B
GFP
24hr-fast cFos
Merge
scramble shRNA
MC3R shRNA
3rd V
C
Num. of cFos positive cells
ad lib fed
24hr fasted
ctrl side
inj side
Scramble shRNA
MC3R shRNA
D
Fold change of cFos positive cell number (norm to ad lib fed)
Scramble shRNA
MC3R shRNA
E
Body weight (g)
Before surgery
After surgery
F
Fat mass (%)
G
Lean mass (%)
scramble
MC3R shRNA
H
WT
Plasma leptin (ng/mL)
Fed
24hr fasted
I
MC3R KO
J
WT
Δ fat mass (g)
Δ plasma leptin (ng/mL)
R2 = 0.4591
p = 0.1392
K
MC3R KO
R2 = 0.4052
p = 0.2482
L
24hr fasting-induced leptin decrement
WT
MC3R KO
% decrease in leptin level (norm to ad lib fed)
M
Plasma Leptin Level
Plasma leptin (ng/mL)
AgRP Cre (+/-)
AgRP Cre (+/-), MC3R (fl/fl)
N
AgRP-Cre+/-
AgRP-Cre+/-, MC3Rfl/fl
Plasma leptin (ng/mL)
Fat mass (g)
R2 = 0.1559
R2 = 0.1471

**Figure S3. Defective AgRP neuron activation in MC3R KO mice is not a result of developmental adaptations or differential reduction in leptin.**

(A) Schematic showing the unilateral injection of scramble or MC3R shRNA virus into the ARC of adult WT male mice.

(B) Representative images showing the expression of scramble or MC3R shRNA and cFos signals in the ARC of adult WT male mice following a 24h fast. Scale bar, 100um.

(C, D) Quantifications of the number of cFos-positive cells and fasting-induced fold change of cFos number in the ARC (n=4~9 mice for all groups).

(E-G) Body weight and body composition of scramble or MC3R shRNA injected WT male mice (n = 8~16 mice for each group).

(H, I and L) Fasting-induced leptin level change in WT and MC3R KO male mice (n = 5~6 mice for each group).

(J, K) Relationship between decreased plasma leptin level and decreased fat mass in response to 24h fasting in WT and MC3R KO male mice.

(M, N) Plasma leptin levels and the relationship between leptin level and fat mass in AgRP-Cre and AgRP-specific MC3R KO male mice (n = 8 mice for each group).

Data are plotted as mean and all error bars represent the SEM. ns, non-significant; *$p < 0.05$; **$p < 0.01$; ***$p < 0.001$, ****$p < 0.0001$ in unpaired Student's t-test and two-way ANOVA with Sidak's posthoc test.
